# Metagenomic analysis of concrete bridge reveals a microbial community dominated by halophilic Bacteria and Archaea

**DOI:** 10.1101/2022.09.16.508313

**Authors:** E. Anders Kiledal, Mark Shaw, Shawn W. Polson, Julia A. Maresca

## Abstract

Concrete hosts a small but diverse microbiome that changes over time. Shotgun metagenomic sequencing would enable assessment of both diversity and function of the microbial community in concrete, but because the biomass in concrete is so low, this analysis is highly affected by laboratory contamination. Here, we demonstrate improved DNA extraction from concrete, and show that this method provides DNA of sufficient quality and quantity to do shotgun metagenomic sequencing. DNA was extracted from a sample of concrete obtained from a road bridge and sequenced with Illumina MiSeq. This microbial community was dominated by halophilic Bacteria and Archaea, with enriched functional pathways related to osmotic stress responses. Prior work found that halophilic bacteria were relatively rare in younger concrete samples, which had abundant oligotrophic taxa. These results suggest that as concrete ages and weathers, salt and osmotic stresses become more important selective pressures, and suggest that long-term persistence and performance of microbes for biorepair or biosensing applications might improve if halophilic strains were used.

## Introduction

Concrete is a dry, salty, oxidized, alkaline environment that hosts a small but diverse community of microbes ^1,2^. These characteristics make it a harsh environment that presumably imposes strong selective pressures on the microbes in it from the time it is poured, but in our previous work, relatively few known halophilic or alkaliphilic taxa were observed during the first two years after curing ^1^. However, it may take years for extreme environments to be colonized by microbes capable of thriving there, and even longer for them to reach detectable levels.

We previously found that in the early months of concrete weathering, microbial communities in concrete resembled those of the precursor materials used in the concrete mix ^1^. Over two years, the diversity of bacteria decreased and became more similar to the microbial communities associated with the large aggregates and cement powder, potentially indicating that these components share characteristics with concrete. We hypothesized that as concrete ages, halo- and alkaliphiles would be more likely to survive than non-extremophilic microbes. To test this hypothesis, we extracted DNA from an 81-year-old bridge in New Jersey and used Illumina MiSeq to generate a shotgun metagenomic data set.

Shotgun metagenomics is more useful than amplicon sequencing for obtaining detailed functional information about microbial communities. However, concrete has such low biomass that metagenomic sequencing poses a real challenge: contaminating DNA from the laboratory and reagents often comprises a relatively large fraction of the extracted DNA and therefore a large number of the sequences obtained. Here, we report an improved method for extracting DNA from concrete samples, adapted from ancient DNA methods ^3,4^. We then extracted and sequenced DNA from a crumbling concrete bridge sample, revealing a microbial community largely composed of halophiles, including many Archaea.

## Results

### Improved DNA extraction from concrete

Several modifications to our DNA extraction protocol were evaluated for their ability to increase DNA yield while limiting background contamination. We previously extracted DNA from concrete samples with a chloroform-based protocol modified from one designed for DNA extraction from ancient bone ^2,5^. The first step of that protocol is a lengthy EDTA wash; longer (48-72 hours) EDTA washes and multiple EDTA washes did not increase DNA yield (Table 1). Tests with the calcium indicator murexide showed that 10 mmol EDTA was sufficient to chelate all of the calcium leached from concrete (data not shown).

**Table 1.**
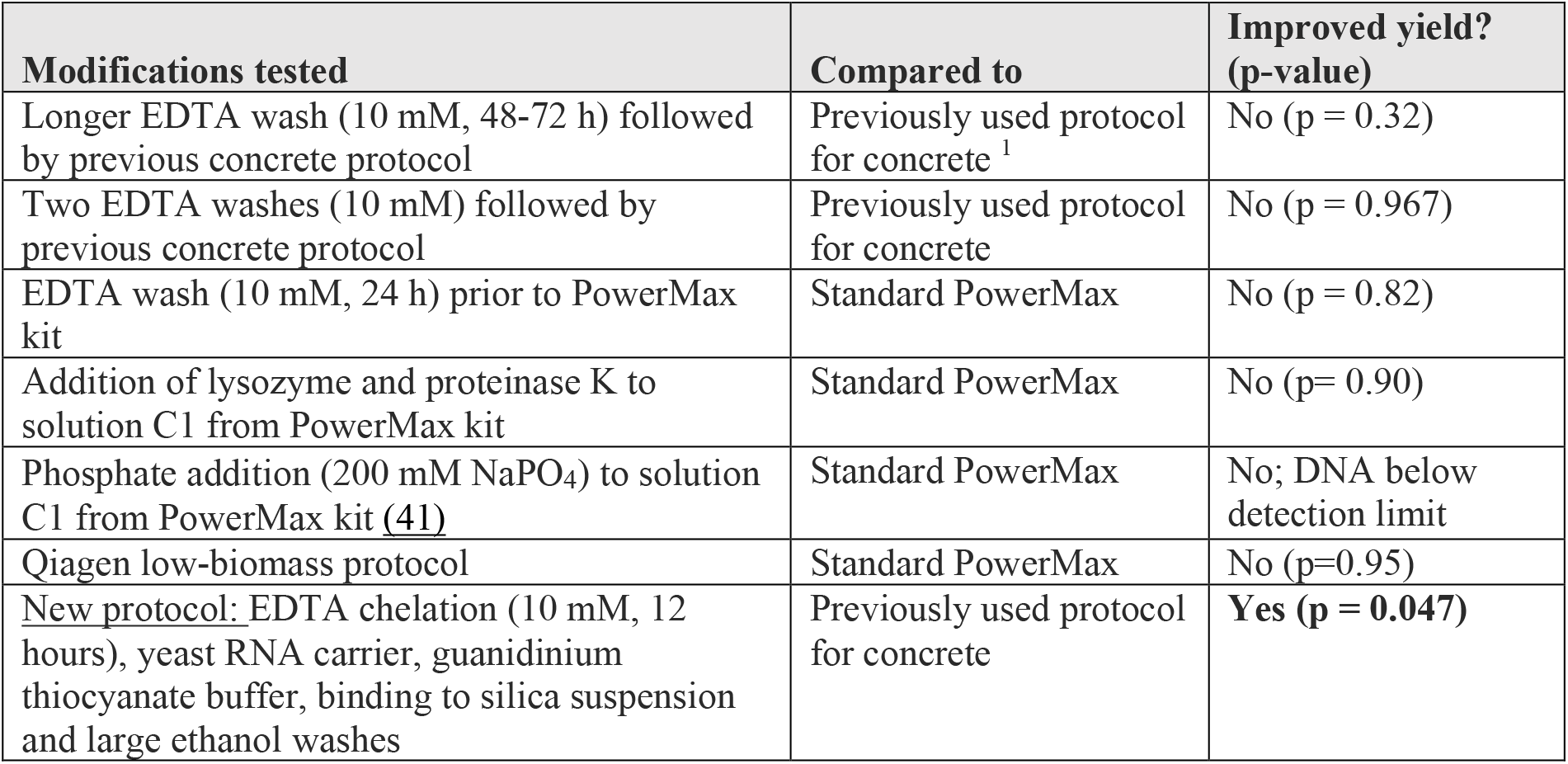
Differences in DNA extraction yields for multiple protocols and protocol modifications. DNA extraction yields were compared between protocols and between modified or unmodified protocols. Improved DNA yield was tested with one-sided Welch’s t-tests; tests with p ≤ 0.05 were considered a significant improvement (bold). No significant protocol modifications were identified. The new concrete DNA extraction protocol using a guanidinium thiocyanate buffer and silica suspension–scaled up from prior ancient DNA extraction techniques–significantly improved DNA yields compared to our previous chloroform-based concrete extraction protocol. The Qubit limit of detection is approximately 0.005 ng µL^-1^.

The Qiagen PowerMAX DNA extraction kit was also used to extract DNA from concrete, with and without modifications. In addition to the standard kit protocol, we tested washing the concrete with EDTA prior to lysis, adding lysozyme and proteinase K to the lysis buffer from ^2^ prior to use of the PowerMax kit, adding proteinase K to the kit lysis buffer (Powerbead solution and solution C1), adding phosphate and ethanol to the kit lysis buffer ^6^, and a low-biomass protocol provided by Qiagen (personal communication). These modifications did not improve yields (Table 1), and some even reduced the amount of DNA recovered. Of the methods evaluated, the PowerMax kit consistently had the smallest yields (Figure 1), with DNA often undetectable, even using a high-sensitivity dsDNA Qubit fluorometric assay with the largest sample volume supported (20 μl).

**Figure 1.**
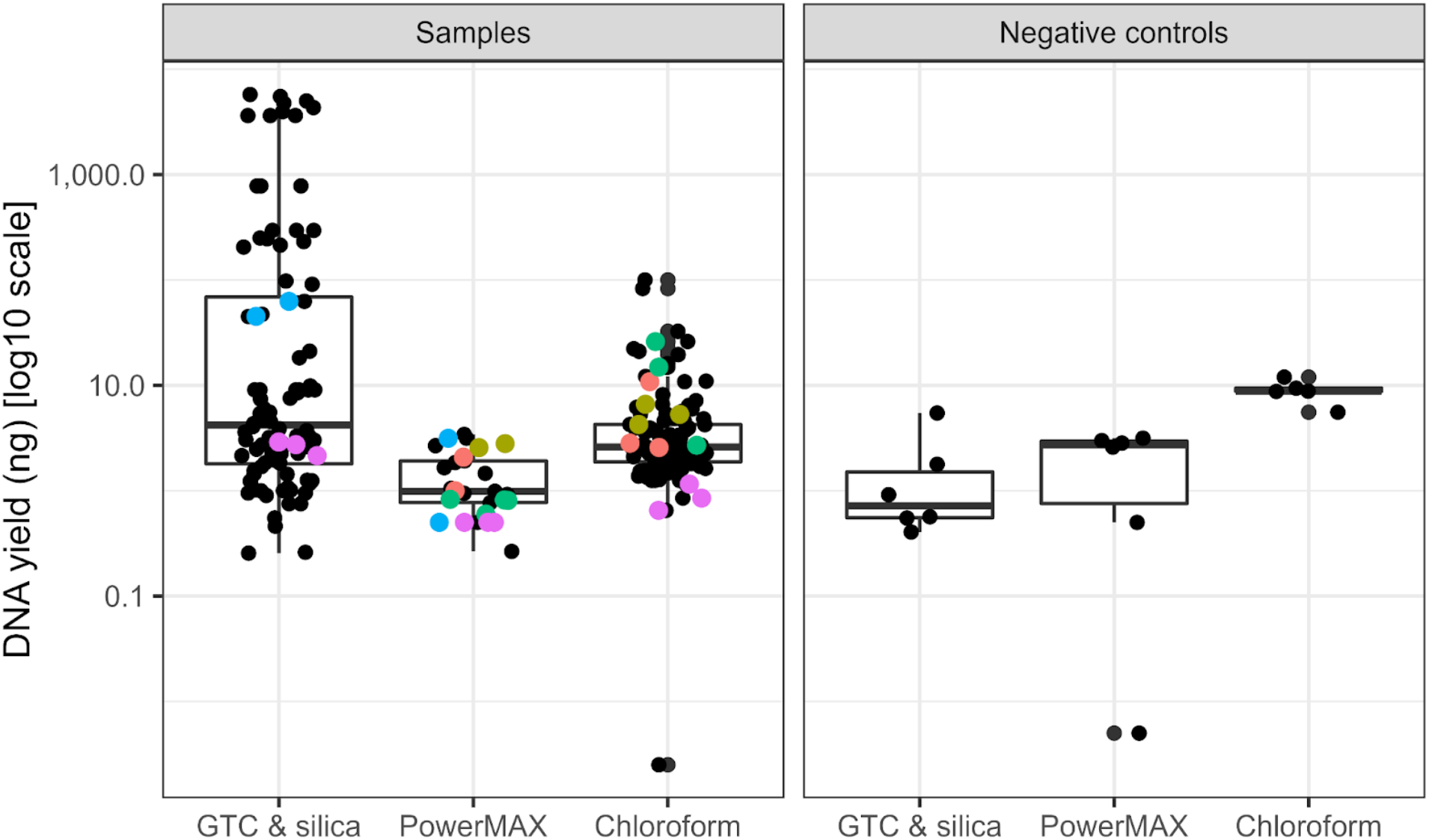
DNA yields using different protocols. Three DNA extraction methods were compared for extracting DNA from concrete: a method that binds DNA to silica particles in a guanidinium thiocyanate buffer (GTC & silica), the Qiagen PowerMAX kit, and our previous chloroform-based extraction protocol chloroform (Chloroform). In each case, DNA was extracted from 10-g concrete samples; yields in ng are presented from both concrete (A) and negative control samples (B). Colors indicate samples extracted with multiple methods. Lower negative control extraction yields were observed with the GTC & silica protocol (one-sided Welch t-test with PowerMAX extractions p = 0.024; one-sided Welch t-test with chloroform extractions p = 0.00021).

Many challenges of extracting DNA from concrete are shared with extracting DNA from ancient bone and teeth samples, and the ancient DNA field more generally ^3,4,7^. These shared challenges include low-biomass samples and the related issue of contamination, and large quantities of divalent cations. A commonly used ancient DNA extraction protocol ^3,4^ was adapted for concrete by scaling up the initial steps to accommodate more material while maintaining small final resuspension volumes to obtain concentrated DNA. The extraction protocol uses an initial EDTA chelation and enzymatic lysis, followed by binding of DNA to a fine silica suspension in a guanidine thiocyanate buffer. Yeast RNA is included as a blocking agent and carrier molecule. This protocol increased DNA yield from concrete samples (Table 1), decreased the yields from blank extractions (Figure 1), is relatively inexpensive, and requires fewer steps than either our previous chloroform-based extraction method or the Qiagen PowerMAX kit.

### Shotgun metagenomic sequencing

Illumina MiSeq sequencing generated 1,240,164 raw read-pairs from the DNA extracted from concrete; no reads were obtained from the negative control extraction. After adapter trimming and quality filtering, 1,237,961 read-pairs remained. The GC content of reads was 61%. Average sequence length before trimming was 154 bp; after trimming the average read lengths were 151 bp (forward) or 150 bp (reverse).

### Community analysis

Reads were first classified with Kraken2 against the Genome Taxonomy Database (GTDB), successfully classifying 767,469 reads (61.99% of the total) as either Bacteria (48.8%) or Archaea (12.8%). Reads unclassified against GTDB (12,155 reads; 0.98% of the total) were further classified with Kraken2 against a RefSeq database including eukaryotes (0.46%) and viruses (0.0036%).

Considering the relative abundance of organisms (percentages that follow are of the 767,469 classified reads) the majority of organisms observed were Bacteria (78.6%), specifically Proteobacteria (42%), Actinobacteria (24%), Firmicutes (7.6%), and Bacteroidota (2%) (Figure 2). Gammaproteobacteria comprised 25% of total observations, particularly halophilic organisms in the Halomonadaceae (12%) which includes *Halomonas*, the most abundant genus observed (9%), *Marinobacter* (1.3%), and *Salinisphaera* (2.3%). Alphaproteobacteria were also abundant (17%), particularly Rhodobacteraceae (6.4%), *Methylobacterium* (2.5%), Rhizobiaceae (2.4%), and Sphingomonadaceae (1.2%). The phylum Actinobacteriota represented 24% of the community, with most belonging to class Actinomycetia (19%). In the Actinobacterial class Rubrobacteria, genus *Rubrobacter* (3.5%) was also abundant. Firmicutes were 7.7% of the community, particularly the genera *Marinococcus* (4.9%) and *Halobacillus* (0.96%).

**Figure 2.**
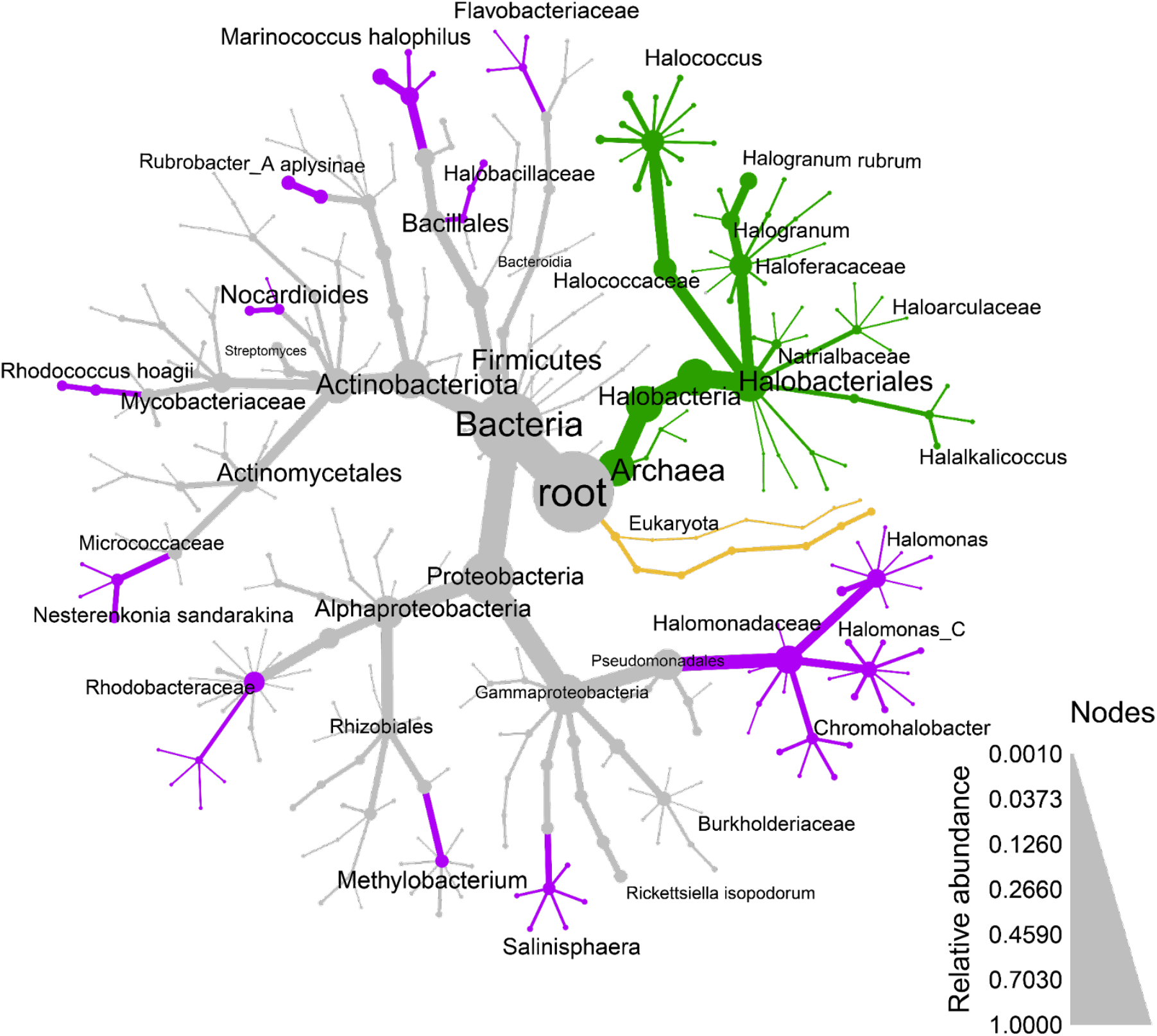
Microbial community composition in concrete. Most of the organisms observed were Bacteria (78.6%) Archaea (20.6%), while eukaryotes and viruses were observed as much lower abundances. All of the observed Archaea are putatively halophilic organisms (green nodes and lines), while many observed Bacteria are also likely halophilic (purple nodes and lines). Node size and edge thickness indicate relative abundance. Only groups with relative abundances > 0.001 are shown; however, excluded taxa were included in abundance calculations of higher-rank taxa.

Archaea (20.6 %), almost exclusively from Halobacteria (20.5%), were abundant in this community and included the species with the highest relative abundance, *Halogranum rubrum* (4.4%; Figure 2). The genus *Halococcus* comprised 7.93% of the community, and we also observed other members of the order Halobacteriales like *Natrialbaceae* (1.41%) and *Halalkalicoccus* (1.2%).

Eukaryotes (0.73%) and viruses (0.0051%) represented only small proportions of the classified reads. The most abundant eukaryotic groups included plants (Streptophyta, 0.56%), and fungi (0.045%). Most metazoan reads (0.12%) were human, likely contamination, while fungi were mostly the plant pathogens *Colletotrichum* and *Fusarium*. Viral sequences observed were mostly bacteriophage of *Mycobacterium, Gordonia, Rhizobium*, or *Acinetobacter*.

### Functional analysis

Enrichment of biological process gene ontology (GO) terms in the concrete metagenomic data relative to all GO annotations of the UniRef90 database was determined by mapping UniRef90 clusters to GO terms and testing for enrichment with the *topGO* R package ^8^ (Table 2). Some of the enriched GO terms include general metabolic strategies such as carbohydrate metabolism, urea catabolism, ethanol oxidation, and photosynthesis. Enriched GO terms related to stress responses include DNA protection ^9^ and heavy metal detoxification, specifically responses to nickel and mercury ions ^10^. Additionally, several pathways related to compatible solutes are enriched, including synthesis and catabolism of ectoine ^11,12^, spermidine and putrescine ^13,14^. Finally, GO terms associated with *Halobacteria* were enriched, including gas vesicle formation ^15^. Complete results of the GO term analysis are available in Supplemental File 3.

**Table 2.**
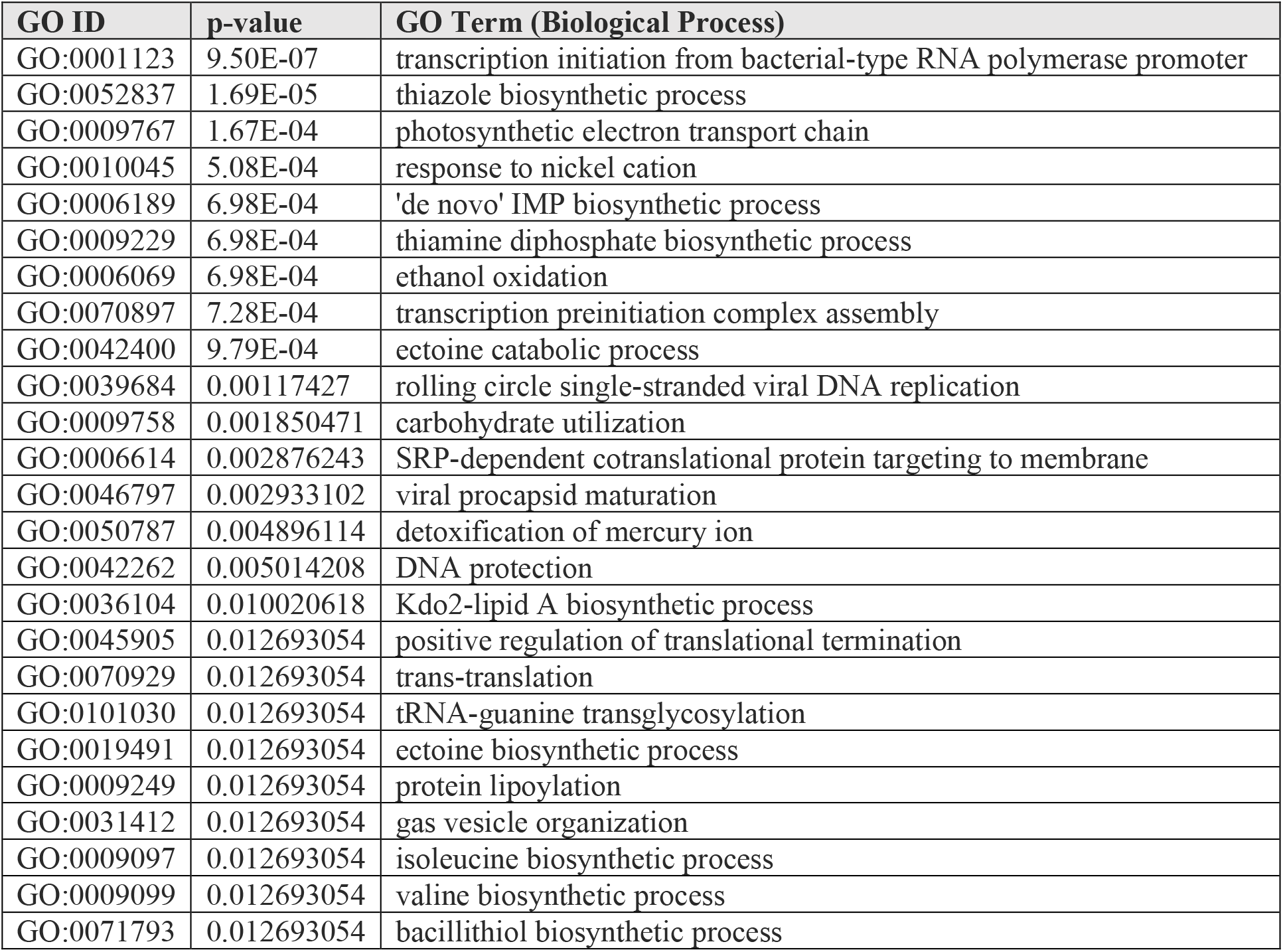
GO Terms Enriched in Metagenomic Data. GO terms enriched in the metagenomic data relative to the UniRef90 cluster GO annotations were identified with the topGO R package, using the weight01 algorithm and Fisher’s exact test (null hypothesis: GO term abundance is the same in the GO term database and concrete metagenomic data). The top 25 most enriched GO terms are included in this table, along with the p-value of the enrichment. A complete list of GO terms and the probabilities that they are enriched in this data set is available in Supplemental File 3.

| GO ID | p-value | GO Term (Biological Process) |
| --- | --- | --- |
| GO:0001123 | 9.50E-07 | transcription initiation from bacterial-type RNA polymerase promoter |
| GO:0052837 | 1.69E-05 | thiazole biosynthetic process |
| GO:0009767 | 1.67E-04 | photosynthetic electron transport chain |
| GO:0010045 | 5.08E-04 | response to nickel cation |
| GO:0006189 | 6.98E-04 | 'de novo' IMP biosynthetic process |
| GO:0009229 | 6.98E-04 | thiamine diphosphate biosynthetic process |
| GO:0006069 | 6.98E-04 | ethanol oxidation |
| GO:0070897 | 7.28E-04 | transcription preinitiation complex assembly |
| GO:0042400 | 9.79E-04 | ectoine catabolic process |
| GO:0039684 | 0.00117427 | rolling circle single-stranded viral DNA replication |
| GO:0009758 | 0.001850471 | carbohydrate utilization |
| GO:0006614 | 0.002876243 | SRP-dependent cotranslational protein targeting to membrane |
| GO:0046797 | 0.002933102 | viral procapsid maturation |
| GO:0050787 | 0.004896114 | detoxification of mercury ion |
| GO:0042262 | 0.005014208 | DNA protection |
| GO:0036104 | 0.010020618 | Kdo2-lipid A biosynthetic process |
| GO:0045905 | 0.012693054 | positive regulation of translational termination |
| GO:0070929 | 0.012693054 | trans-translation |
| GO:0101030 | 0.012693054 | tRNA-guanine transglycosylation |
| GO:0019491 | 0.012693054 | ectoine biosynthetic process |
| GO:0009249 | 0.012693054 | protein lipoylation |
| GO:0031412 | 0.012693054 | gas vesicle organization |
| GO:0009097 | 0.012693054 | isoleucine biosynthetic process |
| GO:0009099 | 0.012693054 | valine biosynthetic process |
| GO:0071793 | 0.012693054 | bacillithiol biosynthetic process |

Ectoine and hydroxyectoine are compatible solutes produced by Bacteria and Archaea in high-salt environments ^12,16,17^. They are synthesized from aspartate by EctBAC(D), and ectoine is imported into cells by the UehABC transporter and degraded by EutDE-Atf-Ssd (Figure 3A) ^12^. Of the (hydroxy)ectoine biosynthesis genes, only EctB was observed in contigs from Archaea (Figure 3B). Homologs of EctCD (the ectoine and hydroxyectoine synthases) were observed in contigs originating from both *Firmicutes* and *Actinobacteria*. The *Gammaproteobacteria* appear to encode all four (hydroxy)ectoine biosynthetic genes, the ectoine transporter, and all of the necessary genes for degradation of ectoine (Figure 3B). The *Halobacteria* appear to encode all of the necessary genes for ectoine degradation.

**Figure 3.**
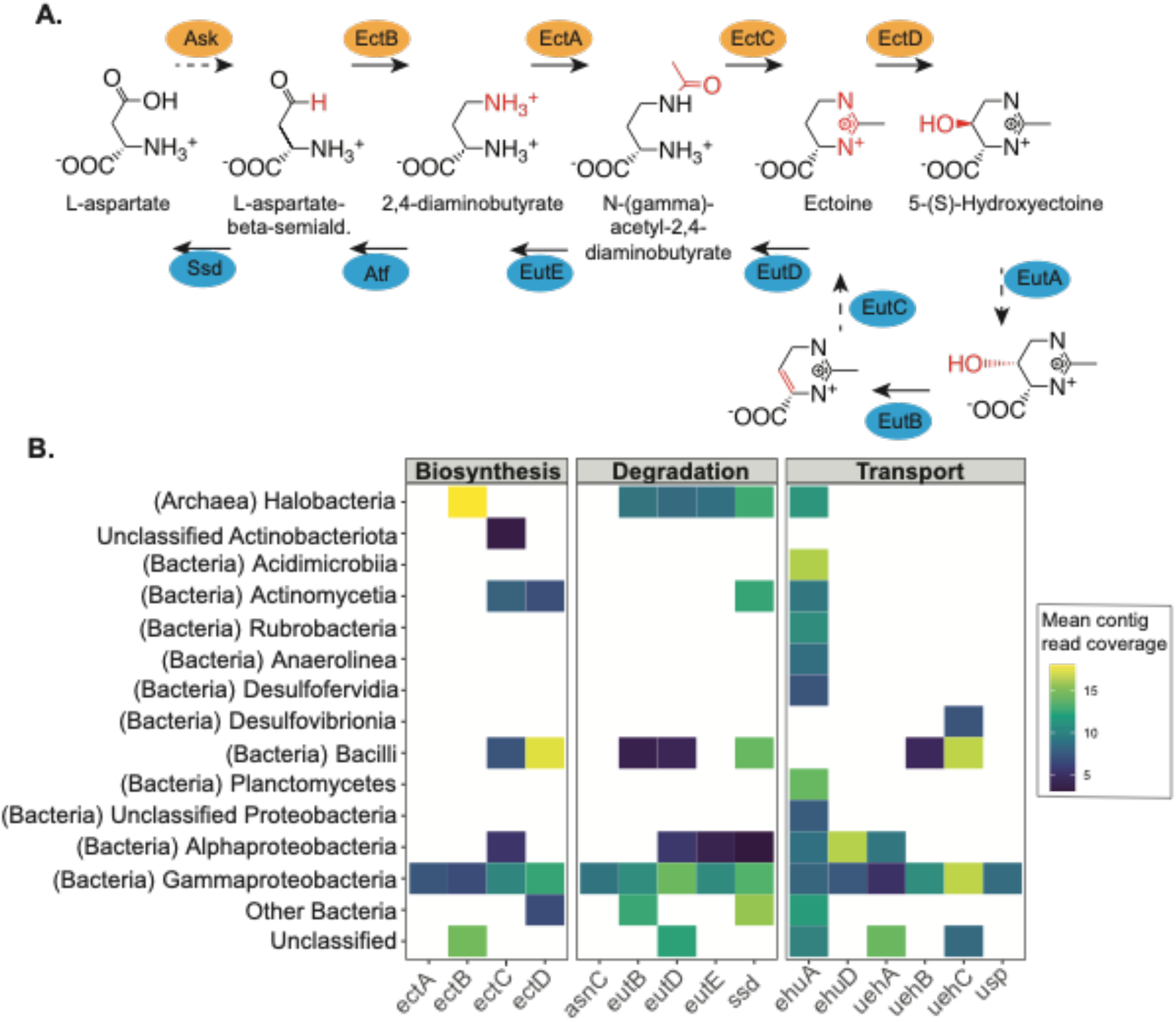
Ectoine synthesis, degradation, and transport. (A) Biosynthesis and degradation pathways for ectoine and hydroxyectoine, after Schulz et al. (B) Genes related to ectoine synthesis, degradation, or transport are found in contigs from a phylogenetically diverse range of microbes, including Halobacteria, several classes of Actinobacteria, and Proteobacteria. Color indicates depth of coverage for the contig with the specified genes.

## Discussion

Here, we analyzed DNA extracted from an 81-year-old, ASR-affected concrete bridge in New Jersey, and found a microbial community dominated by halophiles, with a large number of halophilic Archaea, and very few eukaryotes. These microbes encode pathways that potentially allow them to mitigate the primary selective pressures in concrete, including heavy metal tolerance and a variety of osmotic stress responses. This contrasts with our previous work in much younger concrete (< 3 years old), which used 16S amplicon sequencing to characterize a microbial community composed primarily of generalist, sometimes oligotrophic bacteria, and no detectable Archaea ^1^.

The microbial community in this concrete bridge is highly enriched in halophiles and haloalkilophiles. The differences between this sample and those analyzed in our prior work suggest that the microbial community in concrete can indeed change dramatically with time, and thus that concrete itself is a dynamic (if slow) environment. As concrete ages and reacts with CO_2_ in the atmosphere, the pH decreases to ∼9 ^18^. Further, ASR causes formation of a hygroscopic, alkali-rich gel inside the concrete, and can lead to extensive cracking ^19^. This damage allows water to enter the structure, which likely delivers nutrients to the interior of the concrete, but also solubilizes ions in the concrete matrix. Any microbes able to obtain nutrients from the water would thus also have to tolerate a solution with extremely high salt concentrations and likely high pH ^20^. This strong selective pressure is reflected in the microbial community, which includes not just known halophiles, but also haloalkaliphiles like *Halalkalicoccus* and *Natrialbaceae* ^21,22^.

Although a low-nutrient environment might be expected to select for autotrophic organisms, no carbon fixation pathways are enriched in the metagenome. Because concrete is highly oxidized, there may not be enough reducing power available to support carbon fixation. Even if reduced compounds are released by other organisms in concrete, the biomass in concrete is so low that metabolite transfer may be impossible. One interesting avenue of future research might be to analyze the spatial distribution of microbes throughout concrete.

The differences between the microbial community in this aged, damaged structure and those in young (less than 3 years old) concrete suggests that the microbial communities change over time. The large number of reads attributable to Archaea (∼30%) were a surprise, since no Archaea were detected in any of the samples in our previous work. The primers used in our previous study (357F/806R) do not amplify 16S genes from Halobacteria, so it is unknown whether Archaea were not observed due to primer specificity, or absence/low abundance in the samples. The current results raise the very interesting question of where the microbes in concrete come from. Our prior work indicated that, at least initially, the components of concrete “seed” microbes in concrete. Many of these microbes die within a few months, presumably unable to withstand the conditions in concrete. Did the halophiles in this New Jersey sample descend from a very small number of halophiles present in the initial concrete mix? Alternatively, were they introduced by wind, rain, road salt, or other vectors?

Also somewhat surprising was the low fungal abundance. Fungi, particularly lichens, can colonize concrete surfaces ^23^. It is possible that some of the nearly 40% of reads that could not be classified belong to poorly-characterized fungal groups, since our databases of fungal genomes lag significantly behind those for Bacteria and Archaea ^24^.

This work demonstrates that with an improved, scaled-up method for DNA extraction, shotgun metagenomic analysis of microbial communities within concrete is feasible. This method will facilitate analysis of concrete microbiomes in studies of environmental factors that contribute to infrastructure support or degradation, biorepair of structures by endogenous, bio-augmented, or designed microbial communities, or in efforts to engineer living materials. Here, we show that the microbial community in aging structural concrete is dominated by putatively aerobic, halophilic species, and that the microbial community in old concrete is quite different from microbial communities in relatively new concrete. Perhaps if microbes are added to concrete mixes for future biorepair, a cocktail of strains that will persist and perform at different stages of the concrete service lifetime will be necessary.

## Materials and methods

### Sample collection and processing

The concrete sample was collected from the US 206 bridge over Dry Brook in Sussex County, New Jersey (GPS coordinates 41.140338, −74.743574). To avoid damaging the structure, the sample was collected from a large piece of concrete that had broken off from the underside of the bridge deck. A roughly 500 cm^3^ sample was wrapped in multiple layers of aluminum foil, transported at ambient temperature, and frozen at −80°C the same day. Prior to DNA extraction, concrete fragments were ground to powder in a ring and puck mill that had been scrubbed with soap and hot water, dried, and then cleaned 2x with ethanol and UV irradiated for 40 minutes. A negative control extraction blank was also prepared using 10 g of triple-sterilized glass beads.

### DNA extraction

To increase the DNA yield and decrease contamination introduced during DNA extraction, we tested a number of modifications to the protocol (see Supplemental File 1). After development of the modified protocol, DNA was extracted from 10 g of ground concrete from a New Jersey bridge and a blank negative control extraction. After grinding, all materials were handled inside a UV- and bleach-decontaminated PCR hood. DNA was quantified using an Invitrogen Qubit Fluorometer (version 2) with a Qubit™ dsDNA HS Assay Kit (ThermoFisher catalog #Q32851).

Samples were incubated overnight with gentle rotation at 55°C with 5 ml 0.5M EDTA, 150 µl Proteinase K (800 units ml^-1^; New England Biolabs #P8107S), 138 µl 20% SDS, and 0.2 ml acetic acid in a 50 ml conical tube, scaled up from methods previously described ^3,4,25^. After incubation, samples were vortexed at maximum speed for 10 minutes on a multi-tube vortexer (Dade cat. no. S8215-1). Samples were then centrifuged for 3 minutes at 5000 RPM and the supernatant was transferred to a new 50 ml conical tube. 30 ml of modified QG buffer (5 M guanidium thiocyanate, 18.1 mM Tris-HCl, 25 mM NaCl, 1.3% Triton X-100) was added to the supernatant with 5 µl yeast RNA (1 mg ml^-1^; Sigma-Aldrich #R6625) and 125 µl of a silica suspension prepared as previously described ^25^. DNA was allowed to bind overnight to silica particles at room temperature with gentle mixing (∼30 RPM). Following binding, samples were centrifuged for 5 minutes at 5000 RPM, and the supernatant was discarded. The silica pellet with bound DNA was washed by resuspension in 10 ml 80% ethanol and centrifuged for 10 minutes at 5000 RPM. The supernatant was again discarded and the pellet was resuspended in 1 ml 80% ethanol, transferred to a 1.5 ml microcentrifuge tube, centrifuged for 3 minutes at 13000 RPM, and the supernatant discarded. The silica pellet was air dried in a laminar flow hood. DNA was eluted from the silica twice in 50 µl pre-warmed (60°C) 10 mM Tris for 5 minutes with gentle rotation at 60°C. For each elution, the silica was pelleted for 1 min at 13000 RPM prior to recovery of eluted DNA. This protocol has been published online with additional comments ^26^.

### Library preparation & sequencing

Prior to sequencing library preparation, DNA was quantified with an AATI FEMTO Pulse. Sequencing libraries for the bridge sample and a negative control sample were prepared with the Illumina Nextera XT kit following the manufacturer’s instructions with three modifications: half reactions due to low input DNA, 12.5 minute tagmentation, and 20 PCR cycles. Libraries were quantified with an AATI Fragment Analyzer (fragment length distribution ∼150 bp to 1600 bp) and subsequently size selected using a Sage Scientific BluePippin, resulting in a final library with ∼350 to 650 bp fragments. Libraries were sequenced on an Illumina MiSeq with 150 bp paired-end reads.

### Bioinformatic and statistical analyses

#### Quality control

Reads were filtered and trimmed with *trim galore!* version 6.6 ^27^. *FastQC* version 11.9 ^28^ was used to generate quality reports, which were summarized using Multiqc version 1.6 ^29^. Bioinformatics, data processing, and statistical analyses were conducted with a custom snakemake workflow available on GitHub (https://github.com/MarescaLab/concrete_metagenome_test). Software dependencies in the workflow are managed by Conda and Docker to aid reproducibility.

#### Taxonomic analysis

Kraken2 version 2.1.1 ^30^ was used to generate a taxonomic profile from paired, quality controlled reads. Reads from Bacteria and Archaea were first classified against a Genome Taxonomy Database (GTDB) release 95 database ^31^ produced with Struo ^32^ and available from ftp.tue.mpg.de/ebio/projects/struo/GTDB_release95/. Reads from Eukarya and viruses were classified against a precomputed RefSeq database (PlusPFP 1/27/2021) available at https://benlangmead.github.io/aws-indexes/k2. From Kraken2 taxonomic profiles, Bracken ^33^ was used to calculate species level relative abundance with otherwise default parameters.

#### Assembly

Reads were assembled with MetaSPADES ^34^ version 3.14.1 using default parameters.

#### Functional analysis

Functional annotation was performed with the HUMAnN3 tool ^35^, using a GTDB release 95 ^31^ database produced with STRUO2 ^32,36^ (available from http://ftp.tue.mpg.de/ebio/projects/struo2/) and taxonomic annotations produced with Kraken2 and Bracken ^30,33^. The abundance of UniRef90 gene clusters ^37,38^ was determined using the HUMANN3 tool ^35^ and a mapping file provided by the authors of the HUMAnN3 tool was used to map Gene Ontology (GO) terms to the UniRef90 clusters.

#### Analysis of ectoine-related pathways

Sequences of bacterial and archaeal proteins involved in ectoine biosynthesis, transport, and degradation ^11,12^ (see Supplemental File 2) were used as queries in a TBLASTN (version 2.11.0; ^39^) analysis to identify corresponding genes in the MetaSPADES assembly. Hits were retained if they satisfied the following filtering criteria: E-Value < 0.001, Bit_Score > 60, Percent_ID > 40, percent_of_query_aligned > 20. Contig taxonomy was determined with Kraken2 version 2.1.1 ^30^ and a Genome Taxonomy Database (GTDB) release 95 database ^31^ produced with Struo ^32^ and available from ftp.tue.mpg.de/ebio/projects/struo/GTDB_release95/. Mean contig coverage was determined by mapping reads to contigs with Coverm version 0.6.1 ^40^ and the minimap2 aligner ^41^.

#### Statistical analysis

Enrichment of biological process GO terms was calculated with the topGO R package, using the default *weight01* method which assigns greater weight to more specific GO terms ^8,42^.

## Data Availability

Raw and assembled DNA sequences are available at the NCBI, associated with BioProject <u>PRJNA846790</u>.

## Acknowledgements

This research was supported by the Delaware Department of Transportation with funding also from the Federal Highway Administration (Task 27-1891). E.A.K. was supported in part by the Delaware Environmental Institute Fellows Program, University of Delaware. Support from the University of Delaware Center for Bioinformatics and Computational Biology Core Facility and use of the BIOMIX compute cluster were made possible through funding from Delaware INBRE (NIGMS P20 GM103446), the State of Delaware, and the Delaware Biotechnology Institute.

We also gratefully acknowledge Brewster Kingham and the University of Delaware Sequencing and Genotyping Center for use of the Qubit, sequence library preparation, and the sequence data, and Ryan Rathbun from the New Jersey Department of Transportation for access to the concrete sample. Rock grinder access and training were provided by Deb Jaisi and Ted (Qiang) Li at the University of Delaware.

## Author information

### Author Contributions

EAK did the field collection, DNA extractions, and sequence data analysis, and wrote the first draft of the manuscript. MS advised on strategies for metagenomic sequencing from low-biomass samples, prepared the sequencing libraries, and did the sequencing. SWP advised EAK on the metagenomic analyses. JAM is the principal investigator on this project: she oversaw the laboratory work and data analysis and wrote the final version of the manuscript. All authors approved the final version of the manuscript.

## Supporting Information

**File S1:** Information about additional DNA extraction methods tested but not used (pdf document).

**File S2:** Fasta file of proteins involved in ectoine synthesis, transport, and metabolism, used as queries in BLAST analysis (text file)

**File S3:** Gene Ontology terms and the probability that each one is enriched in this data set (tsv file)

