## Supplemental File 1 for "Metagenomic analysis of concrete bridge reveals a microbial community dominated by halophilic Bacteria and Archaea"

E. Anders Kiledal,[a] Mark Shaw,[b] Shawn W. Polson,[c] Julia A. Maresca[d]#
[a]Department of Biological Sciences, University of Delaware
[b]Sequencing and Genotyping Center, University of Delaware
[c]Center for Bioinformatics and Computational Biology, University of Delaware
[d]Department of Civil and Environmental Engineering, University of Delaware

**Supplemental methods: Modification of DNA extraction protocol**

Modifications to the DNA extraction method were tested on subsamples from concrete cylinders described previously [1], concrete samples obtained from New Jersey bridges, and concrete samples obtained from a former factory pad on the campus of the University of Delaware. For all DNA extraction experiments, DNA was quantified using an Invitrogen Qubit Fluorometer (version 2) with a Qubit™ dsDNA HS Assay Kit (ThermoFisher catalog #Q32851).

To increase the DNA yield and decrease contamination introduced during DNA extraction, we tested a number of modifications to the protocol. To ensure that calcium and other divalent cations were removed from solution, samples were incubated in EDTA (10 mM) for 12, 48, and 72 hours at 22°C prior to cell lysis. A murexide titration [2] was used to confirm that the amount of EDTA added was sufficient to chelate all $Ca^{2+}$ released from the concrete. Briefly, 5 g of concrete was incubated overnight with shaking in 100 mL sterile distilled water to allow $Ca^{2+}$ to fully dissolve. Murexide powder and NaCl were mixed in a 1:200 (w/w) ratio and 0.5 grams of this mixture was added to the concrete solution. EDTA was titrated into the concrete solution until the color changed to violet; 5 mL of 0.5 M EDTA was required to fully chelate the $Ca^{2+}$.

Because the biomass in concrete is low, we tested DNA extraction protocols developed for other low-biomass environments. Qiagen provided a modified protocol to use with the DNEasy PowerSoil and DNEasy Soil PowerMax kits (Cat. nos. 12888-50 and 12988-10, respectively) to extract DNA from low-biomass samples.

Yeast RNA (Sigma-Aldrich cat. no. R6625) was added to the extraction buffer to block potential nucleic acid binding sites in the concrete matrix. To confirm that the RNA did not add contaminating DNA, a concentration gradient of yeast RNA (0.5 to 20 µg RNA per reaction) was tested for DNA first by quantifying DNA in the solution with the HS DNA Qubit kit, then by using yeast RNA as the templates in PCR reactions using the 16S primers 357F/806R (reaction conditions 95°C for 5 min., 35 × [95°C/30 sec., 47.5°C/30 sec., 72°C/1 min.]). No PCR products were detected (Figure S1), and double-stranded DNA was not detectable by Qubit (limit of detection ~0.005 ng $\mu L^{-1}$) in the RNA solution.

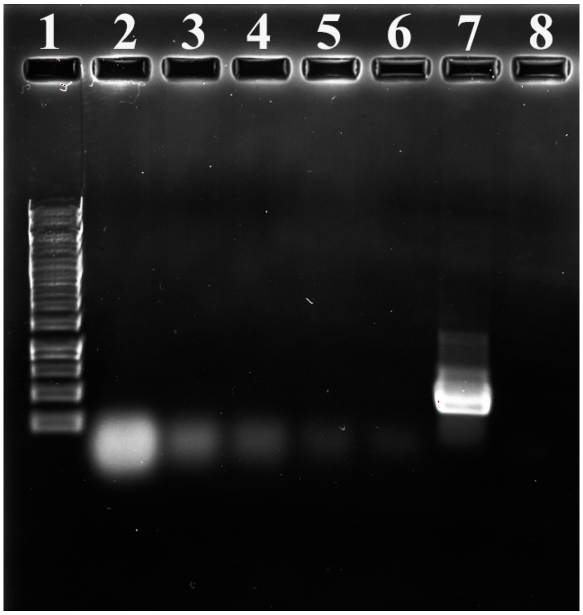


**Figure S1. Yeast RNA solution has no observed bacterial 16S PCR amplification products.**

Several dilutions of the yeast RNA solution used as a carrier molecule in DNA extractions from concrete were tested for bacterial DNA contamination by PCR amplification with the universal bacterial 16S rRNA gene primers 357F and 806R. PCR volumes were 25 μl in all cases. No detectable amplification was observed, indicating that the yeast RNA is not contaminated with bacterial DNA. Lanes: (1) 1kb ladder (ThermoFisher), (2) PCR reaction with 20 μg yeast RNA, (3) PCR reaction with 10 μg yeast RNA, (4) PCR reaction with 5 μg yeast RNA, (5) PCR reaction with 1 μg yeast RNA, (6) PCR reaction with 0.5 μg yeast RNA, (7) PCR reaction with *E. coli* DNA (positive control), (8) PCR with no template DNA (negative control).
